# Sexual dimorphism of autism-like phenotypes in microglial eIF4E overexpression mice

**DOI:** 10.64898/2026.09.03.748596

**Authors:** Changran Niu, Juan Ji An, Hannah V. Masterson, Baoji Xu

**Affiliations:** Department of Neuroscience, The Herbert Wertheim UF Scripps Institute for Biomedical Innovation & Technology, University of Florida, Jupiter, FL 33458, USA; Skaggs Graduate School of Chemical and Biological Sciences, The Scripps Research Institute, La Jolla, CA 92037, USA

**Keywords:** Six keywords: autism spectrum disorder, microglia, dendritic spine, sexual dimorphism, sex hormone, four-core-genotype mice

## Abstract

**Background:** Autism spectrum disorder (ASD) is a group of neurodevelopmental disorders characterized by deficits in social communication and interaction, and restricted interests or repetitive behaviors. ASD is approximately four times more prevalent in males than in females. In this study, we investigated whether sex hormones or sex chromosomes underlie the male bias in ASD susceptibility.

**Methods:** We used the MG^4E^ mouse model, in which microglial eIF4E overexpression produces robust male-biased ASD-like phenotypes. To distinguish the contributions of sex hormones and sex chromosomes, MG^4E^ mice were crossed with four-core-genotype (FCG) mice carrying *Sry* gene manipulations, generating eight genotypes of experimental mice. Social interaction and repetitive behaviors were assessed using standard behavioral assays. Dendritic spine density was quantified in Thy1-GFP mice. Expression of estrogen receptors (ERs) and androgen receptor (AR) was examined in microglia isolated from control and MG^4E^ mice.

**Results:** Sex hormones, rather than non-*Sry* genes on sex chromosomes, are responsible for the male-biased deficits in social interaction and the increase in dendritic spine density. At postnatal day 14, ERs were undetectable in microglia, whereas ER expression was readily detected in neurons.

**Conclusions:** These findings demonstrate that sex hormones are a major determinant of the increased susceptibility of males to ASD-like phenotypes in MG^4E^ mice. The absence of ER expression in microglia suggests that sex hormones may act indirectly through hormone-responsive neurons to regulate microglia-neuron interactions during neurodevelopment.

## Introduction

Autism spectrum disorder (ASD) is a heterogenous neurodevelopmental disorder characterized by two core diagnostic features: deficits in social communication, and restricted interests or repetitive behaviors. The prevalence of ASD has increased steadily over the past two decades, with the current estimate from the Centers for Disease Control and Prevention indicating that 1 in 31 children aged 8 years in the United States is diagnosed with ASD (1). Notably, ASD is diagnosed approximately four times more frequently in males than in females, yet the biological mechanisms underlying this pronounced sex bias remain poorly understood (1, 2).

ASD is a multifactorial disorder with a strong genetic component. Over 1,000 genes have been implicated is ASD and are cataloged in the Simons Foundation Autism Research Initiative (SFARI) Gene database. Despite this remarkable genetic heterogeneity, individuals with ASD share common behavioral features, suggesting that diverse genetic defects may converge on a limited number of pathogenic mechanisms. Dysregulation of translational control has emerged as one such convergent pathway (3, 4). Loss-of-function mutations in several translation repressors, including TSC1, TSC2, PTEN and FMRP, are associated with ASD in humans (5–7). Eukaryotic translation initiation factor 4E (eIF4E), which acts downstream of these ASD-associated translation regulators, controls the translation of a subset of mRNAs with highly structured, G/C-rich 5′ untranslated regions (UTRs) (8–10). Mutations and single nucleotide polymorphisms (SNPs) in the *EIF4E* gene have also been associated with ASD (11–13).

Furthermore, mouse models with 4E-BP deficiency or eIF4E overexpression exhibit excitation/inhibition (E/I) imbalance and ASD-like behaviors (14, 15). Together, these findings underscore the importance of eIF4E-dependent translational control in ASD pathogenesis.

Because there are multiple cell types in the brain, identifying the cellular populations in which dysregulated translational control contributes to ASD is critical for understanding disease mechanisms. Increasing evidence suggests that microglia may represent on such key cellular target. Microglia are resident immune cells in the central nervous system (CNS) that continuously survey the brain microenvironment and respond to developmental and pathological cues. During brain development, microglia play essential roles in establishing neural circuits by regulating axon guidance, neurite growth, synapse formation, and synaptic pruning (16, 17).

Growing evidence implicates microglial dysfunction in ASD. Postmortem studies have reported increased microglial activation and density in individuals with ASD (18–21). In addition, aberrant microglia-mediated synaptic pruning has been observed in several mouse models of neurodevelopmental disorders, including ASD (*Scn2a*-deficient mice) (22), fragile X syndrome (*Fmr1* knockout mice) (23), and Rett syndrome (*Mecp2* null mice) (24). These findings support an important role for microglial dysfunction in ASD pathogenesis.

We previously demonstrated that microglia-specific eIF4E overexpression (MG^4E^) is sufficient to induce ASD-like phenotypes, including increased spine density, E/I imbalance, and impaired sociability (25). Remarkably, these phenotypes were observed only in male MG^4E^ mice, making this model one of the few that recapitulates the striking male bias characteristic of human ASD. The two principal biological factors distinguishing males and females are gonadal hormones and sex chromosome complement (26). The *Sry* gene on the Y chromosome encodes the testis-determining factor required for the formation of testes in males (26). During the perinatal development, males experience a surge of testicular testosterone, much of which locally converted to estradiol in the brain, whereas females are not exposed to comparable levels of gonadal hormones until puberty (26). Consequently, sexually dimorphic hormone exposure during early development may contribute to the male-biased ASD-like phenotypes observed in MG^4E^ mice. In addition, females possess two X chromosomes, whereas males possess one X and one Y chromosome. Although X-inactivation largely equalizes gene dosage between the sexes, a subset of X-linked genes escape inactivation, resulting in sex-specific differences in gene expression (27). Such differences in sex chromosome complement may also contribute to the sex bias in ASD susceptibility.

In this study, we combined the MG^4E^ mouse model and the Four Core Genotype (FCG) mouse model, which dissociates the effects of gonadal sex from those of sex chromosome complement through manipulation of the *Sry* gene. This strategy enabled us to determine whether the sexually dimorphic ASD-like phenotypes in MG^4E^ mice are driven primarily by gonadal hormones specified by the *Sry* gene or by non-*Sry* genes on the sex chromosomes. By separating these two major determinants of biological sex, we sought to define the mechanism underlying the male-biased susceptibility to ASD-like phenotypes in this model.

## Methods and Materials

### Animals

Mouse strains, XY^-^Tg(Sry) [B6.Cg-Tg(Sry)2Ei *Sry^dl1Rlb^* T(XTmsb4x-Hccs;Y)1Dto/ArnoJ; stock #010905] and Thy1-GFP [Tg(Thy1-EGFP)MJrs/J; stock #007788], were obtained from the Jackson Laboratory. The MG^4E^ mice were generated in our lab as previously described (25). Both male and female mice were used in the experiments. All experiments were performed in accordance with relevant guidelines and regulations regarding the use of experimental animals. The Animal Care and Use Committees at the UF Scripps Institute approved all animal procedures used in this study (protocol 16-003-04).

### Tamoxifen treatment

Tamoxifen (Sigma #T5648-5G) was dissolved in 100% ethanol at a concentration of 20 mg/ml. An equal volume of corn oil was then added, and the mixture was vortexed thoroughly. After centrifugation at 13,000 rpm for 10 min, the solution was placed in a vacuum centrifuge for 30 mins to evaporate the ethanol. Tamoxifen was administered to neonatal mice at postnatal day 0 (P0) by subcutaneous injection into the dorsal neck region at a dose of 180 mg/kg body weight to induce nuclear translocation of the CreER fusion protein.

### Behavioral assays

#### Open field test

Locomotor activity and anxiety-like behavior were assessed using the open field test. Each mouse was placed in the center of an open field arena (43.8 x 43.8 x 32.8 cm) and allowed to explore freely for 30 min. Total distance traveled and time spent in the center zone were recorded automatically using the EthoVision XT video tracking system (Noldus Information Technology). The arena was cleaned with 0.5% Quatricide between trials to eliminate olfactory cues. Behavioral analyses were performed by an experimenter blinded to genotype.

#### Three-chamber sociability test

Sociability was assessed using a three-chamber apparatus consisting of three interconnected compartments. A test mouse was first placed in the center chamber and allowed to freely explore the empty apparatus for 10 min (habituation phase).

Immediately afterward, an unfamiliar stimulus mouse of the same sex and a similar age was placed inside a wire mesh enclosure in one side chamber, while an identical empty enclosure was placed in the opposite side chamber. The location of the stimulus mouse was randomized between experiments. The test mouse was then allowed to explore all three chambers for 10 min. Time spent in each chamber was recorded using the EthoVision XT video tracking system. The apparatus and wire mesh enclosures were cleaned with 0.5% Quatricide between trials to eliminate olfactory cues. Behavioral analyses were performed by an experimenter blinded to genotype.

### Dendritic spine analysis

For dendritic spine analysis, z-stack images of secondary dendrites were acquired at 0.2-μm intervals using an Olympus FV3000 laser-scanning confocal microscope equipped with a 60x oil-immersion objective (NA 1.50, WD 0.11 mm). Dendrites were traced on maximum-intensity projection images, and their lengths were measured using the SNT plugin (Neuroanatomy) in FIJI. Dendritic spines were manually counted using the Point Tool in FIJI. Spine density was calculated as the number of dendritic spines per micrometer of dendrite length. All image acquisition and quantitative analyses were performed by an experimenter blinded to genotype.

### Immunohistochemistry

Mice were transcardially perfused with phosphate-buffered saline (PBS), followed by 4% paraformaldehyde (PFA) in PBS. Brains were dissected and postfixed in 4% PFA at 4°C for 24 h, followed by cryoprotection in 30% sucrose in PBS. Coronal brain sections (40 μm thick) were cut using a sliding microtome (Leica).

Free-floating sections were incubated in 10% methanol and 3% hydrogen peroxide in Tris-buffered saline (TBS) to quench endogenous peroxidase activity, and then blocked in TBS containing 0.3% Triton X-100 and 10% normal horse serum. Sections were incubated overnight at 4°C with primary antibodies diluted in blocking buffer. The following primary antibodies were used: guinea pig anti-Iba1 (1:500; Synaptic System, #234308) and rabbit anti-ERα (1:1000; Invitrogen, #PA1-309). After three washes with blocking buffer, sections were incubated with the appropriate horseradish peroxidase-conjugated secondary antibody for 1 h at room temperature. For ERα staining, followed washes with TNT buffer, immunoreactivity was visualized using TSA-FITC (Akoya Biosciences, #NEL701A001KT). Sections were then counterstained with DAPI, mounted onto glass slides, and coverslipped with Fluoromount-G containing DAPI (SothernBiotech, #0100-20). Images were acquired using a Nikon C2+ confocal microscope.

### Microglial isolation

Microglia were isolated from the whole brains of P14 mice. Brains were minced into approximately 1-mm^3^ pieces in dissection medium and incubated in papain solution containing 30 U papain (Worthington, #LS003126) and 150 U DNase I (Sigma-Aldrich, #DN25) at 37°C for 1 h. The tissue was then transferred to 15-ml tubes containing Earle’s Balanced Salt Solution (EBSS) supplemented with 0.5% bovine serum albumin (BSA) and mechanically dissociated by gentle trituration until a homogenous cell suspension was obtained. The suspension was filtered through a 40-μm cell strainer and centrifuged at 300x g for 8 min at 4°C. The supernatant was discarded, and the cell pellet was resuspended in ovomucoid protease inhibitor solution (Worthington, #LK003182). Microglia were subsequently isolated by magnetic-activated cell sorting using CD11b MicroBeads (Miltenyi Biotec, #130-093-634) according to the manufacturer’s instructions.

### Quantitative RT-PCR

Freshly isolated microglia were resuspended in RNAlater (Sigma-Aldrich, #R0901) and stored at -80°C until RNA extraction. Total RNA was extracted using the Absolutely RNA Nanoprep Kit (Agilent, #400753) according to the manufacturer’s instructions. First-strand cDNA was synthesized from purified RNA using SuperScript IV Reverse Transcriptase (Invitrogen, #18090050) with random hexamer primers. Quantitative PCR (qPCR) was performed using SYBR Green 2x Master Mix (Applied Biosystems, #A25742) in a total reaction volume of 20 μl on a QuantStudio 6 Pro Real-Time PCR System (Applied Biosystems). Relative gene expression was calculated using the 2^−ΔΔCt method and normalized to the housekeeping gene *Actb*.

## Results

### Generation of eight genotypes of mice

We utilized the Four Core Genotypes (FCG) mouse model, in which XY^-^Tg(*Sry*) males carry a deletion of the endogenous *Sry* gene on the Y chromosome (Y^-^) and a transgenic copy of *Sry* inserted on an autosome (28, 29). XY^-^Tg(*Sry*) mice were crossed with *Rosa^Eif4e/Eif4e^* control (CON) mice for two generations to generate male XY^-^Tg(*Sry*) CON mice (Fig. 1). Male XY^-^ Tg(*Sry*) CON mice were then crossed with female *Rosa^Eif4e/Eif4e^*;*Cx3cr1^CreER^*^/+^ (MG^4E^) mice to generate CON and MG^4E^ offspring representing the four core genotypes: XX, XY^-^, XXTg(*Sry*), and XY^-^Tg(*Sry*) (Fig. 1). In this model, XX and XY^-^Tg(*Sry*) mice correspond to wild-type (WT) females and males, respectively, whereas XXTg(*Sry*) mice are chromosomal females with testes (gonadal males) and XY^-^ mice are chromosomal males with ovaries (gonadal females). All offspring at P0 received tamoxifen (180 mg/kg) to induce CreER-mediated recombination. MG^4E^ mice harbor the microglia-specific Cre driver *Cx3cr1^CreER^*, which activates eIF4E overexpression from the *Rosa^Eif4e^* allele following tamoxifen administration (25).

**Figure 1.**
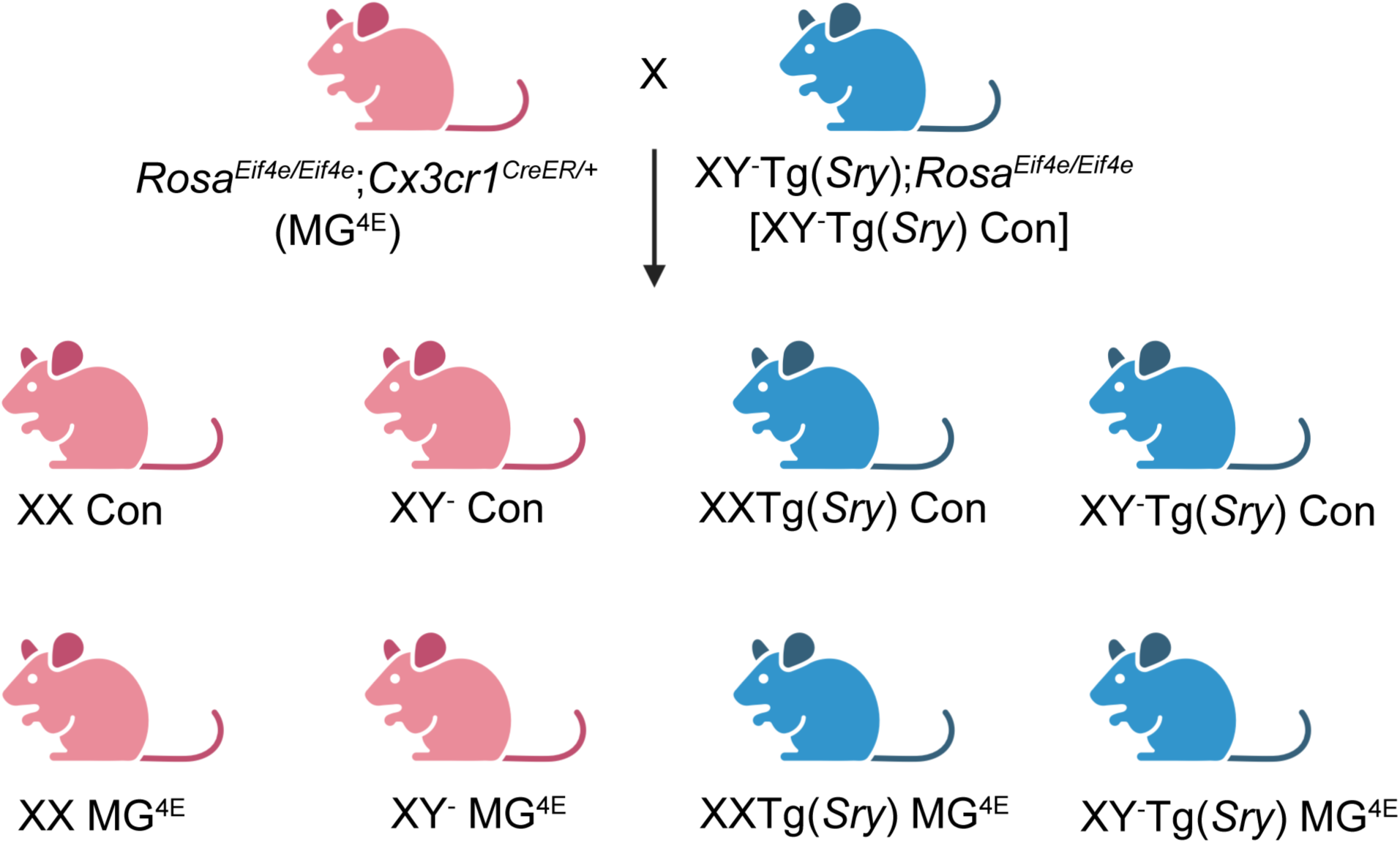
Breeding scheme of generating eight genotypes of mice. XX and XY^-^ mice are phenotypic females. XXTg(*Sry*) and XY^-^Tg(*Sry*) are phenotypic males.

Based on our previous findings, we expected ASD-like phenotypes to develop in XY^-^Tg(*Sry*) MG^4E^ mice but not in XX MG^4E^ mice (25). Comparison of the four MG^4E^ genotypes allowed us to distinguish the effects of gonadal sex from those of sex chromosome complement. Specifically, the presence of ASD-like phenotypes in XXTg(*Sry*) MG^4E^ mice would indicate that gonadal sex is sufficient to confer susceptibility, whereas the presence of ASD-like phenotypes in XY^-^ MG^4E^ mice would instead implicate non-*Sry* genes on the sex chromosomes.

### Sex hormones underlie the male-biased social impairment in MG^4E^ mice

We first examined behavioral phenotypes in the eight experimental genotypes. Open field test (OFT) and three-chamber sociability test were performed at 2-3 months of age. In the OFT, MG^4E^ and CON mice within each FCG group traveled comparable distances, indicating that microglial eIF4E overexpression did not impair locomotor activity (Fig. 2A). To assess anxiety-like behavior, we separately analyzed the first 5 min of the 30-min OFT. MG^4E^ and CON mice spent similar amounts of time in the center zone, indicating that MG^4E^ mice did not exhibit increased anxiety in any FCG group (Fig. 2B).

**Figure 2.**
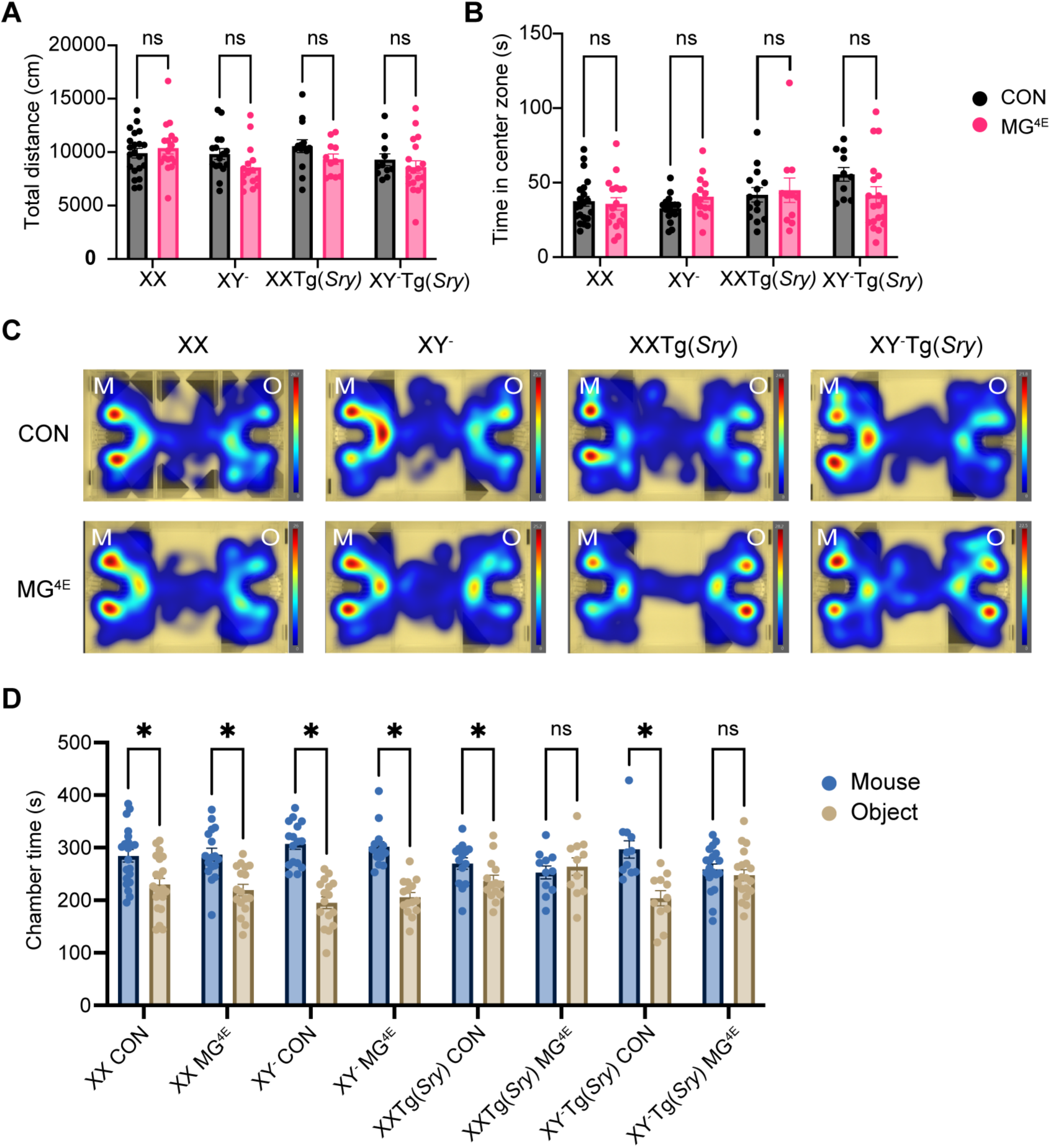
Gonadal sex determines the male-biased deficit in social interaction in MG^4E^ mice. **(A)** Total distance traveled during the 30-min OFT. **(B)** Time spent in the center zone during the first 5 min of OFT. **(C)** Representative heatmaps showing chamber preference during the three-chamber sociability test. M: mouse chamber; O: object chamber. **(D)** Time spent in the mouse and object chambers during the three-chamber sociability test. Sample sizes: XX, n = 21 CON and 17 MG^4E^; XY^-^, n = 16 CON and 15 MG^4E^; XXTg(*Sry*), n = 14 CON and 11 MG^4E^; XY^-^Tg(*Sry*), n = 11 CON and 19 MG^4E^. Two-sided unpaired Student’s *t* test, *p≤0.05; ns, not significant.

We next assessed sociability using the three-chamber test. All CON groups spent significantly more time in the chamber containing a novel mouse than in the chamber containing a novel object, demonstrating normal sociability (Fig. 2C, D). As expected, XY^-^ Tg(*Sry*) MG^4E^ mice failed to show this preference and spent comparable amounts of time in the mouse and object chambers, indicating impaired social interaction (Fig. 2C, D). In contrast, XX MG^4E^ mice retained a significant preference for the mouse chamber, consistent with our previous finding that social deficits in MG^4E^ mice are male biased (25). Importantly, XXTg(*Sry*) MG^4E^ mice also exhibited impaired sociability, whereas XY^-^ MG^4E^ mice displayed normal social interaction, closely resembling XX MG^4E^ mice (Fig. 2C, D). These findings indicate that gonadal sex, rather than non-*Sry* genes on the sex chromosomes, is the primary determinant of male-biased social impairment in MG^4E^ mice. Together, these results demonstrate that the sexual dimorphism in social behavior in MG^4E^ mice is determined predominantly by the sex hormones.

### Sex hormones underlie the male-biased increase in dendritic spine density in MG^4E^ mice

Altered dendritic spine density is a common neuropathological feature observed in ASD (30–32). To determine whether the male-biased increase in dendritic spine density in MG^4E^ mice is governed by gonadal sex or sex chromosome complement, we crossed XY^-^Tg(*Sry*) MG^4E^ with Thy1-GFP mice to sparsely label CA1 pyramidal neurons in the eight experimental genotypes.

Consistent with our previous findings (25), dendritic spine density was significantly increased in XY^-^ Tg(*Sry*) MG^4E^ mice but not in XX MG^4E^ mice compared to their respective control groups (Fig. 3A, B). Importantly, XXTg(*Sry*) MG^4E^ mice also exhibited significantly higher dendritic spine density than XXTg(*Sry*) CON mice, whereas spine density was comparable between XY^-^ MG^4E^ and XY^-^ control mice (Fig. 3A, B). These findings indicate that gonadal sex, rather than non-*Sry* genes on the sex chromosomes, is the primary determinant of the male-biased increase in dendritic spine density in MG^4E^ mice. This conclusion is consistent with the behavioral results, which likewise demonstrate that gonadal sex determines the male-biased social impairment in MG^4E^ mice (Fig. 2C, D).

**Figure 3.**
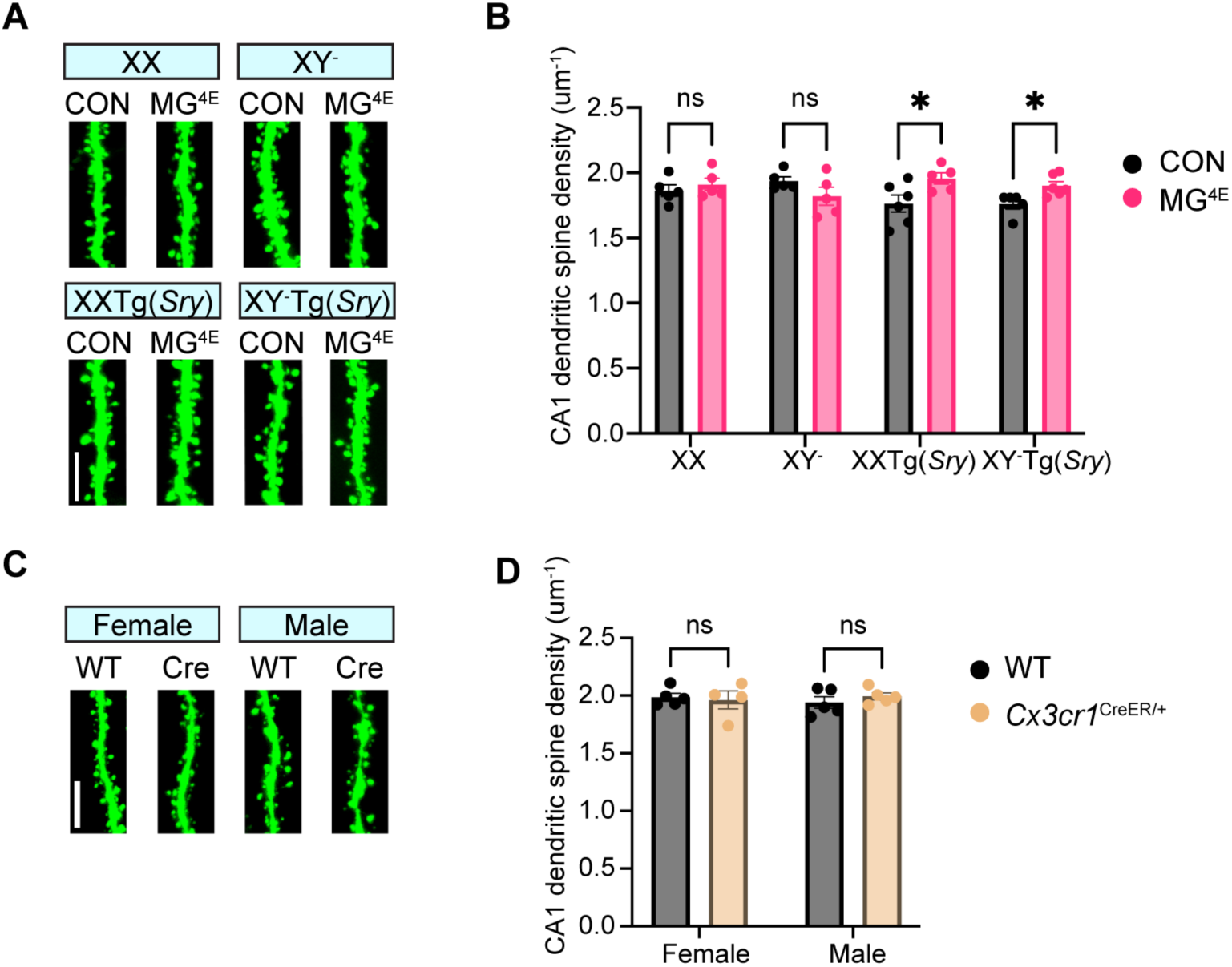
Gonadal sex determines the male-biased increase in dendritic spine density in MG^4E^ mice. (A) Representative images and (B) quantification of dendritic spines on GFP-labeled CA1 pyramidal neurons from 6-week-old Thy1-GFP mice. Sample sizes: XX, n = 5 CON and 5 MG^4E^ mice; XY^-^, n = 5 CON and 5 MG^4E^ mice; XX Tg(*Sry*), n = 6 CON and 5 MG^4E^ mice; XY^-^ Tg(*Sry*), n = 5 CON and 6 MG^4E^ mice. Four to ten dendritic segments were analyzed per mouse. (C) Representative images and (D) quantification of dendritic spine density in CA1 pyramidal neurons from 6-week-old WT and *Cx3cr1^CreER/+^* mice. Female: 5 WT and 4 *Cx3cr1^CreER/+^* mice; Male: 5 WT and 5 *Cx3cr1^CreER/+^* mice. Four to ten dendritic segments were analyzed per mouse. Scale bars, 5 μm. Two-sided unpaired Student’s *t* test: *p≤0.05; ns, not significant.

To exclude the possibility that Cre expression contributed to the increase in dendritic spine density, we compared WT and *Cx3cr1^CreER/+^* mice and found no significant difference between the two groups (Fig. 3C, D). These results indicate that the male-biased increase in dendritic spine density is attributable to microglial eIF4E overexpression rather than the mutation in the *Cx3cr1* allele (Fig. 3C-D).

### The effects of sex hormones are not mediated by microglial ERα

Having established that gonadal sex underlies the male-biased social impairment and increased dendritic spine density in MG^4E^ mice, we next investigated the underlying molecular mechanism. Male mice experience a perinatal surge in testosterone, whereas females do not (26). In the brain, testosterone can signal directly through the androgen receptor (AR) or be locally aromatized to estradiol, which acts through estrogen receptors (ERs) (33). The translation initiation factor eIF4E selectively enhances translation of mRNAs containing highly structured, G/C-rich 5′UTRs (8–10) or cytosine-enriched regulator of translation (CERT) motifs (34).

Interestingly, eIF4E overexpression increases ERα expression in tamoxifen-resistant breast cancer cells (35). We therefore hypothesized that microglial eIF4E overexpression increases the expression of ERs or AR in microglia, thereby enhancing sex hormone signaling during postnatal brain development. Enhanced hormone signaling could disrupt microglial function, leading to impaired synaptic pruning, increased dendritic spine density, and deficits in social behavior.

To test this hypothesis, we first examined the expression of ERs and AR in microglia at postnatal day 14 (P14), when microglial abnormalities are already detectable in MG^4E^ mice (25). Microglia were isolated from P14 CON and MG^4E^ mice, and the purity of the isolated cells was assessed by quantitative RT-PCR. Compared with whole hippocampal tissue, the microglial markers *Cd68* and *Cx3cr1* are highly enriched, whereas the neuronal marker *Syn1* and the astrocytic marker *Gfap* were markedly depleted (Fig 4A), confirming the high purity of the isolated microglia.

**Figure 4.**
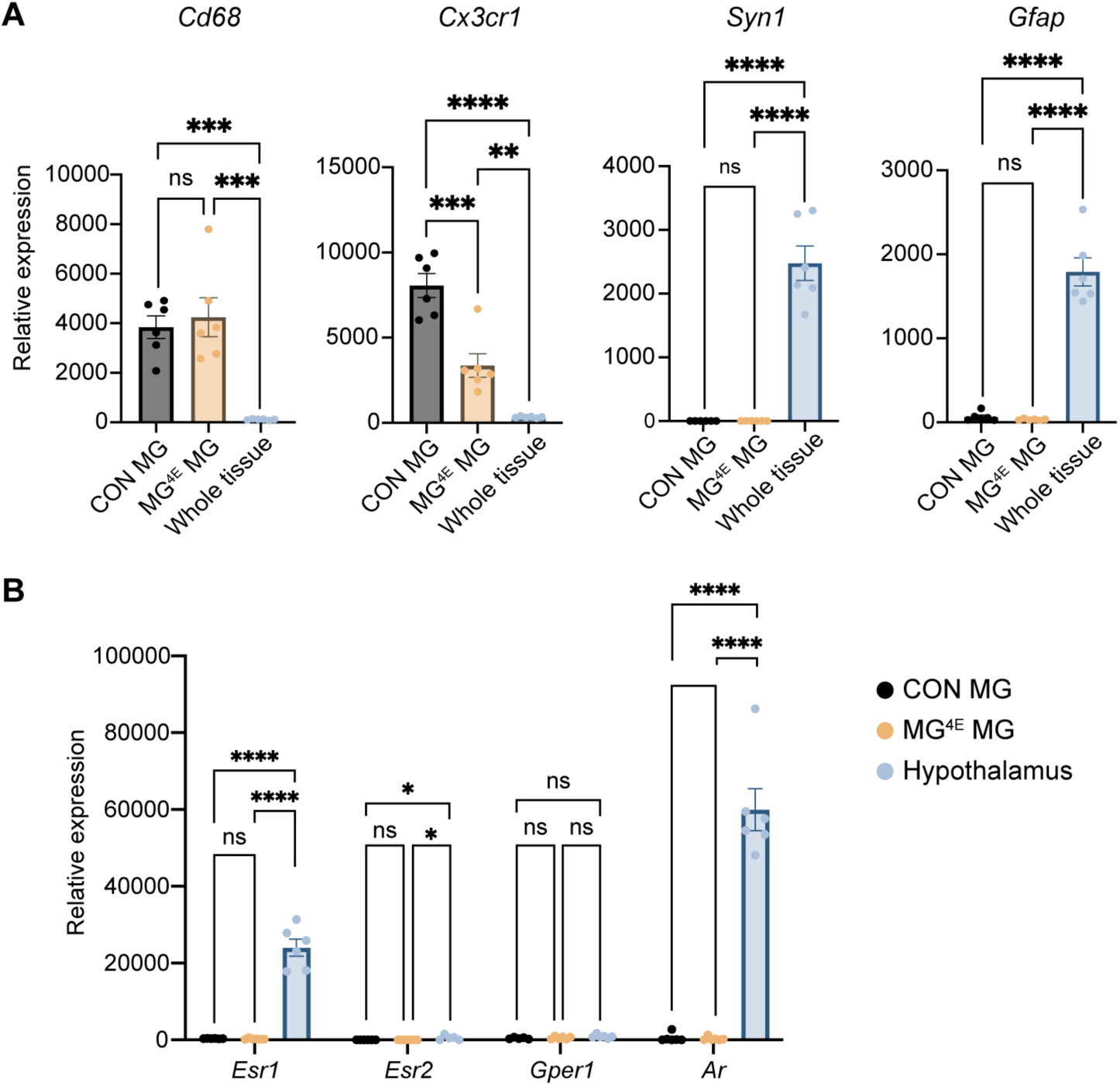
ER and AR mRNAs are undetectable in P14 microglia. (A) Relative expression of the microglial markers *Cd68* and *Cx3cr1* and the neuronal and astrocytic markers *Syn1* and *Gfap* in microglia isolated from P14 mouse brains. Whole hippocampal tissue served as the reference control, confirming the high purity of isolated microglia. (B) Relative expression of *Esr1*, *Esr2*, *Gper1*, and *Ar* in microglia isolated from P14 mouse brains. Hypothelamic tissue served as a positive control for estrogen receptor and androgen receptor expression. Gene expression was normalized to *Actb*. Data are presented as mean ± SEM. One-way ANOVA followed by Tukey’s multiple comparisons test: *p<0.05; **p<0.01; ***p<0.001; ****p<0.0001; ns, not significant.

We next measured the expression of the three estrogen receptors, *Esr1* (ERα), *Esr2* (ERβ), and *Gper1* (G protein-coupled estrogen receptor), as well as *Ar* in isolated microglia. Hypothalamic tissue, which expresses high levels of ERs and AR (36), was used as a positive control.

Expression of *Esr1*, *Esr2*, *Gper1*, and *Ar* was extremely low in both CON and MG^4E^ microglia compared with hypothalamic tissue (Fig. 4B), indicating that microglia express little, if any, ERs or AR at P14.

To further validate the qPCR result, we performed immunohistochemical analysis of ERα in P14 mouse brains. Consistent with published in situ hybridization data (36), ERα immunoreactivity was abundant in the amygdala but relatively low in the hippocampal CA1 region (Fig. 5).

**Figure 5.**
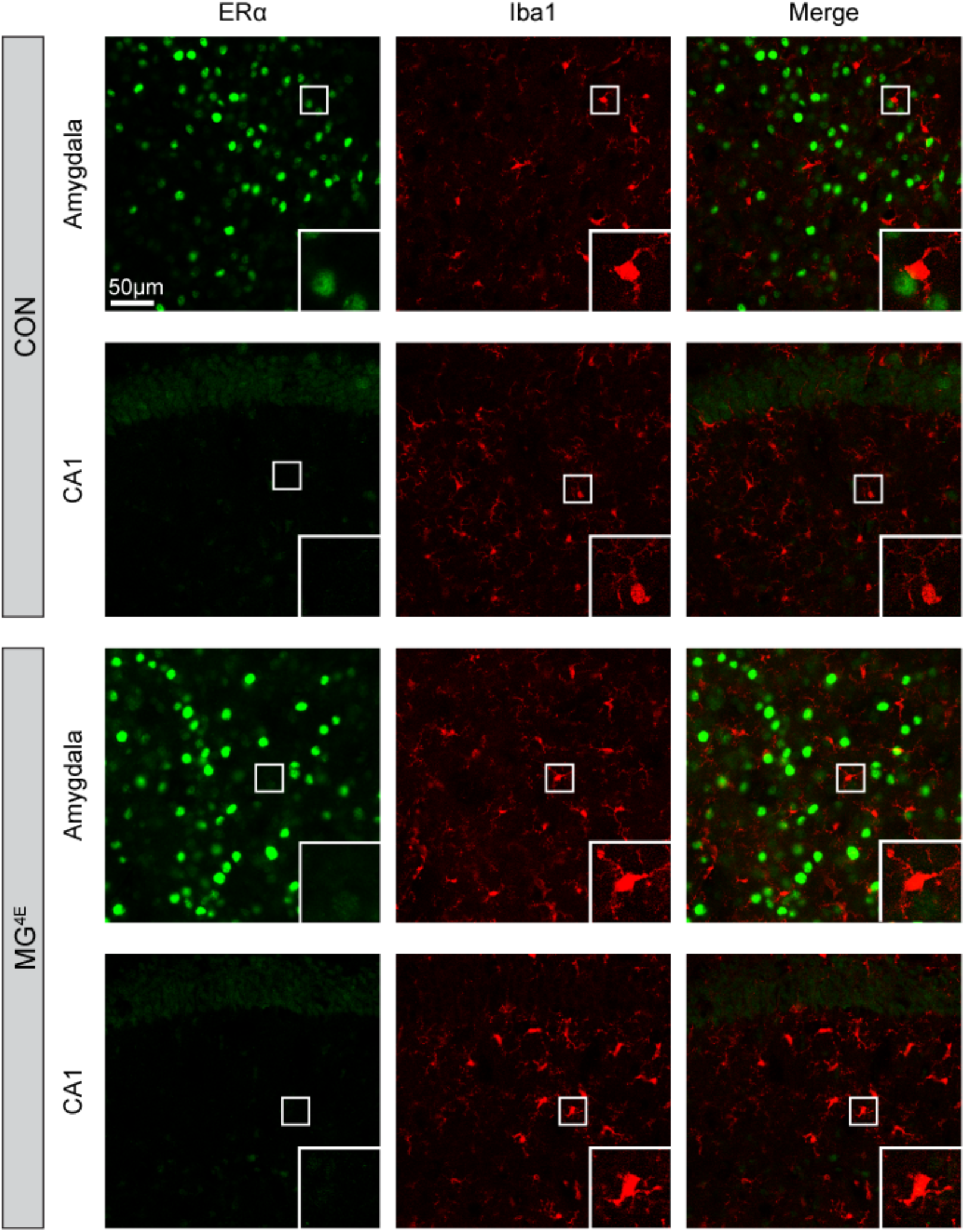
ERα is not detected in microglia at P14. Representative immunofluorescence images of ERα (green) and the microglial marker Iba1 (red) in the amygdala and hippocampal CA1 region of P14 CON and MG^4E^ mice. Insets show 3x magnified views of the boxed regions. No colocalization of ERα and Iba1 was observed in either brain region. Scale bar, 50 μm.

However, ERα immunoreactivity did not colocalize with the microglial marker Iba1 in either the amygdala or CA1 of CON or MG^4E^ mice, indicating that microglia exhibit no detectable ERα expression in these brain regions at p14 (Fig. 5).

Together with the qPCR results, these findings indicate that sex hormone receptors are expressed at little, if any, detectable levels in microglia during postnatal development. Therefore, the male-biased phenotypes observed in MG^4E^ mice are unlikely to result from direct ER- or AR-mediated signaling within microglia. Instead, because ERs and AR are predominantly expressed in neurons, we propose that sex hormones act primarily on neuronal ERs and/or AR, thereby altering communication between neurons and microglia and ultimately disrupting microglial regulation of synaptic pruning (Fig. 6).

**Figure 6.**
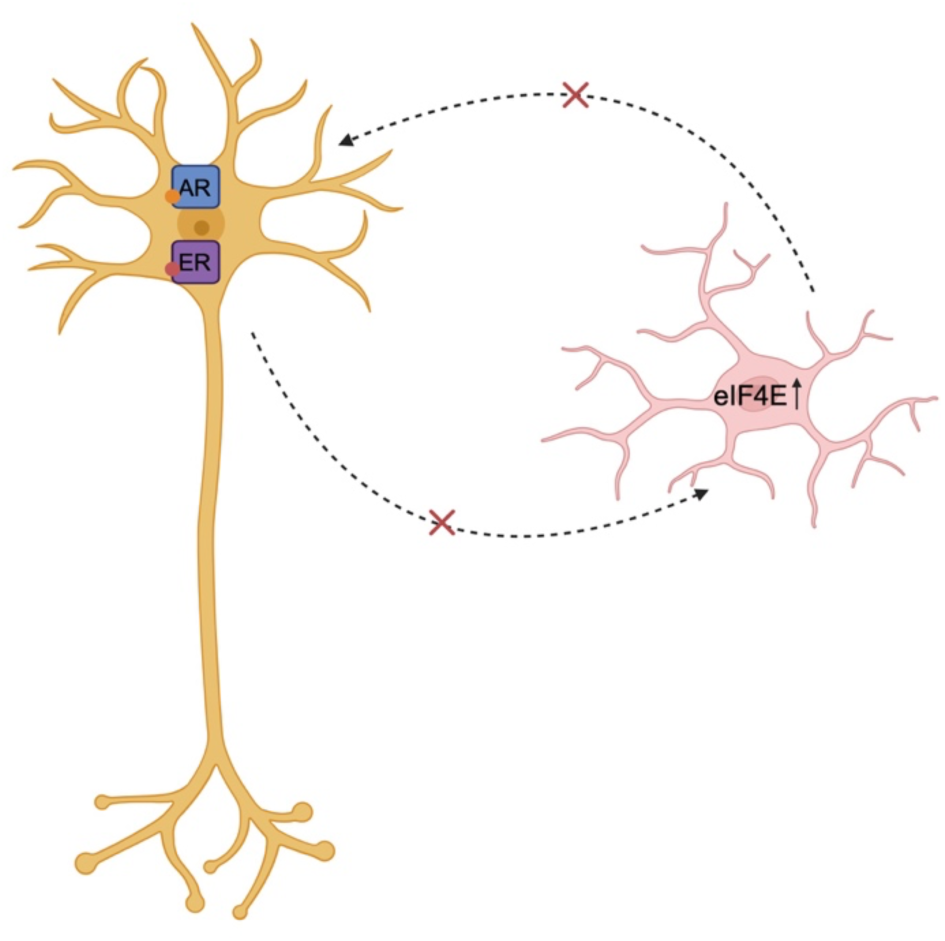
Proposed mechanisms underlying the male-biased phenotypes in MG^4E^ male mice. Sex hormones are proposed to act primarily through neuronal ERs and/or AR, rather than directly on microglia. In combination with microglial eIF4E overexpression, neuronal sex hormone signaling may disrupt communication between neurons and microglia, impair microglia-mediated synaptic pruning, and ultimately increase dendritic spine density and cause deficits in social interaction in male MG^4E^ mice.

## Discussion

Epidemiological studies consistently demonstrate a male predominance in ASD, although the magnitude of this sex difference has decreased in recent years. According to the Autism and Developmental Disabilities Monitoring (ADDM) Network, the male-to-female ratio among 8-year-old children diagnosed with ASD declined from 4.5:1 in 2012 (37) to 3.4:1 in 2022 (1), likely reflecting improved recognition and diagnosis of ASD in females. Nevertheless, females continue to be diagnosed later than males and are more likely to be misdiagnosed because they often present with subtler or atypical clinical features (38, 39). Differences in case ascertainment also influence the reported sex ratio (40). Despite these diagnostic and methodological factors, a male predominance of approximately 3:1 persistent (1, 40), indicating that biological mechanisms substantially contribute to the increased susceptibility of males to ASD.

In this study, we investigated the biological basis of the male predominance using the MG^4E^ mouse model, which uniquely recapitulates both ASD-like molecular and behavioral phenotypes together with a marked male bias. By combining this model with the FCG mouse model, we demonstrated that gonadal sex, rather than non-*Sry* genes on the sex chromosomes, determines the male-biased deficits in sociability and increased dendritic spine density. Because the principal biological consequence of *Sry* expression is the establishment of testes and the resulting perinatal exposure to gonadal hormones, these findings strongly implicate developmental sex hormone exposure as a major determinant of male vulnerability in this model.

Our findings are consistent with clinical studies linking androgen signaling to ASD susceptibility. Elevated fetal testosterone levels have been associated with increased autistic traits (41–43), and conditions associated with increased prenatal androgen exposure, such as maternal stress and polycystic ovary syndrome, have been linked to an elevated risk of ASD (44, 45). Several studies have also reported elevated circulating androgen levels in individuals with ASD (46, 47). However, causal relationships are difficult to establish in humans because of genetic heterogeneity, environmental influences, and ethical limitations. The MG^4E^ mouse therefore provide a valuable experimental model for investigating the mechanisms by which developmental sex hormones influence ASD susceptibility.

Testosterone readily crosses the blood-brain barrier and can be converted to either 17β- estradiol (E2), which signals through ERs, or dihydrotestosterone (DHT), which selectively activates the AR. Although the classic view of brain sexual differentiation emphasizes estrogen-mediated masculinization, accumulating evidence indicates that androgen-AR signaling also contributes independently to this process (48, 49). Determining the relative contributions of ER- and AR-dependent signaling to the male-biased phenotypes observed in MG^4E^ mice will therefore be an important direction for future investigation.

At the cellular level, our findings suggest that sex hormones do not act directly on microglia, as estrogen and androgen receptor expression was minimal in microglia at P14. Instead, the effects of sex hormones are likely mediated indirectly through neurons, where these receptors are abundantly expressed. Our data therefore support a model in which neuronal sex hormone signaling alters neuron-to-microglia communication, ultimately disrupting microglial regulation of synaptic development. Importantly, sex hormones alone are unlikely to be sufficient to produce ASD-like phenotypes. Rather, they appear to interact with underlying cellular vulnerabilities, represented in our model by microglial dysfunction caused by eIF4E overexpression. Consistent with this interpretation, the normal perinatal testosterone surge did not produce ASD-like phenotypes in wild-type mice but, in the presence of microglial eIF4E overexpression, was associated with robust behavioral deficits and increased dendritic spine density in males. These findings support a gene-by-sex interaction model in which developmental sex hormone signaling and microglial dysfunction converge to increase susceptibility to ASD-like phenotypes.

Both AR and ERs can signal through genomic and non-genomic pathways. In the classical genomic pathway, androgen or estrogen binding induces receptor dimerization and nuclear translocation of AR, ERα, or ERβ, allowing these ligand-activated transcription factors to bind androgen or estrogen response elements or interact with other transcription factors to regulate gene transcription (50, 51). In parallel, membrane-associated AR and ERs mediate rapid non-genomic signaling by activating intracellular pathways, including phospholipase C/protein kinase C, Ras/Raf/MAPK, phosphatidyl inositol 3 kinase/Akt, and cAMP/protein kinase A signaling (51, 52). We propose that sex hormones increase susceptibility to ASD-like phenotypes by modulating neuron-microglia interactions in MG^4E^ mice. Sex hormones are well known to induce dendritic spine formation and synaptic plasticity in multiple brain regions (53–55). Through genomic signaling, they may alter the neuronal expression of molecules involved in neuron– microglia communication, including complement proteins, chemokines, cytokines, purinergic signaling molecules, or other synaptic "eat-me" signals that regulate microglia-mediated synaptic pruning (56, 57). Through non-genomic signaling, they may enhance neuronal excitability and intracellular Ca²⁺ signaling, increasing ATP release and other activity-dependent signals that recruit microglia. Because MG^4E^ microglia exhibit impaired motility in response to ATP stimulation (25), these neuronal changes may further compromise microglia-mediated synaptic remodeling, ultimately resulting in increased dendritic spine density and ASD-like behavioral deficits.

In conclusion, this study provides mechanistic insight into the biological basis of the male predominance in ASD by demonstrating that developmental sex hormone signaling interacts with microglial dysfunction to drive male-biased phenotypes. Our findings further suggest that these effects are mediated indirectly through neuronal ER and/or AR signaling rather than through direct hormone action on microglia. Elucidating the molecular pathways that regulate neuron-microglia communication during this critical developmental period may provide new insights into the pathogenesis of ASD and identify novel therapeutic targets for neurodevelopmental disorders.

## Acknowledgements

This work was supported by a grant the National Institutes of Health, United States to B.X. (R01 MH125187). ChatGPT was used to refine human-written text.

## Author contributions

CN and BX designed research; CN, JJA, and HVM performed experiments; CN analyzed data; and CN and BX wrote the paper.

